# Coronary artery disease risk and lipidomic profiles are similar in familial and population-ascertained hyperlipidemias

**DOI:** 10.1101/321752

**Authors:** Joel T. Rämö, Pietari Ripatti, Rubina Tabassum, Sanni Söderlund, Niina Matikainen, Mathias J. Gerl, Christian Klose, athan O. Stitziel, Aki S. Havulinna, Veikko Salomaa, Nelson B. Freimer, Matti Jauhiainen, Aarno Palotie, Marja-Riitta Taskinen, Kai Simons, Samuli Ripatti

## Abstract

**Aims**: To characterize and compare coronary artery disease (CAD) risk and detailed lipidomic profiles of individuals with familial and population-ascertained hyperlipidemias.

**Methods and Results**: We determined incident CAD risk for 760 members of 66 hyperlipidemic families (≥ 2 first degree relatives with the same hyperlipidemia) and 19,644 Finnish FINRISK population study participants. We also quantified 151 lipid species in plasma or serum samples from 550 members of 73 hyperlipidemic pedigrees and 897 FINRISK participants using a mass spectrometric shotgun lipidomics platform. Hyperlipidemias (LDL-C or triacylglycerides over 90^th^ population percentile) were associated with increased CAD risk (high LDL-C: HR 1.74, 95% CI 1.48–2.04; high triacylglycerides: HR 1.38, 95% CI 1.09–1.74) and the risk estimates were very similar between the family and population samples. High LDL-C was associated with altered levels of 105 lipid species in families (p-value range 0.033–7.3*10^−20^ at 5% false discovery rate) and 51 species in the population samples (p-value range 0.017–6.8*10^−21^). Hypertriglyceridemia was associated with altered levels of 117 lipid species in families (p-value range 0.035–1.8*10^−49^) and 119 species in the population sample (p-value range 0.038–2.3*10^−56^). The lipidomics profiles of hyperlipidemias were highly similar in families and population samples.

**Conclusion**: We identified distinct lipidomic profiles associated with high LDL-C and triacylglyceride levels. CAD risk, lipidomic profiles and genetic profiles are highly similar between familial and population-ascertained hyperlipidemias, providing evidence of similar and overlapping underlying mechanisms. Our results do not support different screening and treatment for such hyperlipidemias.

## Introduction

High levels of low-density lipoprotein cholesterol (LDL-C) and triacylglycerides (TGs) have been identified as causal risk factors for atherosclerotic cardiovascular disease (ASCVD).^1, 2^ These hyperlipidemias may arise through lifestyle factors, but they are also highly heritable.^3–6^ More than half of patients with premature coronary artery disease (CAD) may be affected by familial dyslipidemias, a majority of which are characterized by elevations in LDL-C and/or TGs.^7^

Whether familial hyperlipidemias should be diagnosed and managed differently from hyperlipidemias observed in randomly ascertained individuals in the general population is uncertain. Carriers of rare high-impact variants predisposing to familial hypercholesterolemia (FH) may have a higher risk of developing coronary artery disease than non-carriers with similar lipid levels.^8^ This is potentially related to lifelong exposure to high LDL-C levels and suggests that these individuals may benefit from earlier or more aggressive LDL-C-lowering therapy.

Rare variants, however, explain only a small fraction of familial hyperlipidemias with either elevated LDL-C or TG levels. Instead, there is growing evidence that familial hyperlipidemias are highly polygenic in origin.^8–13^ It is of high clinical importance whether non-monogenic familial hyperlipidemias represent distinct disease entities and lead to different ASCVD susceptibility compared with population ascertained hyperlipidemias.^14^ A recent study provided suggestive evidence to the contrary, finding greater preclinical atherosclerosis in monogenic FH than in polygenic hypercholesterolemia.^15^ However, it is unknown how these results translate to clinical outcomes between familial and population-ascertained hyperlipidemias.

How the heterogenic genetic background of familial hyperlipidemias manifests in detailed circulating lipid profiles, and reflects underlying pathophysiology, is also unknown. Recent technological advancements have allowed replicable and simultaneous quantification of hundreds of lipid species through lipidomic profiling.^16, 17^ We tested whether precise phenotypic differences in lipidomic profiles, which might underlie differing ASCVD susceptibility, exist between familial and population-ascertained hyperlipidemias after excluding individuals with monogenic FH. We looked beyond the traditional clinical measures of LDL-C and TG concentrations at the numerous fatty acid ester species of TGs and cholesterol species which are the major constituents of LDL and TG-rich lipoprotein particles. We used a direct infusion platform that combines absolute quantification with high throughput, overcoming problems that have hampered many previous studies.^18^

Lipid species including sphingolipids, glycerophospholipids, glycerolipids and cholesteryl esters may predict ASCVD incidence or event risk over traditional risk factors, and the levels of many individual lipid species have been linked to distinct genetic loci.^19–22^ Major differences in the metabolic pathways underlying different types of hyperlipidemias would thus be expected to be reflected in different lipidomic profiles. As an example, individuals with low HDL-C levels have previously been shown to have low phosphatidylethanolamine-plasmalogen levels in HDL particles, a marker of HDL anti-oxidative capacity.^23^

In the present study, we first estimated the CAD risk associated with familial and population ascertained hyperlipidemias. Secondly, we characterized the lipidomic profiles associated with elevated plasma levels of LDL-C and TGs. Finally, we compared the lipidomic profiles of familial and population-ascertained hyperlipidemias to assess their potential differences.

## Materials and Methods

### Subjects and clinical ascertainment

The Finnish hyperlipidemia families included in this cohort study (74 families, *n* = 1,445 individuals with at least LDL-C and TG measures) were identified as part of The European Multicenter Study on Familial Dyslipidemias in Patients with Premature Coronary Heart Disease (EUFAM). Designation of “familial high LDL-C” or “familial high TGs” was made if at least two first-degree relatives of each other had LDL-C or TG levels, respectively, that were > 90^th^ age and sex-specific Finnish 1997 population percentiles (Supplemental Table IV). Classic familial hypercholesterolemia (FH) was excluded based on genotyping and an in-house functional low-density lipoprotein receptor test.

Samples from the Finnish National FINRISK study were used as a Finnish population-based comparison group. 19,644 individuals from the FINRISK 1992–2002 cohorts and 755 individuals from EUFAM families passed exclusion criteria and could be linked with the national hospital discharge and causes-of-death registries. Clinical incident CAD event endpoints were defined as either myocardial infarction or coronary revascularization (coronary angioplasty or coronary artery bypass grafting). Mean (range) follow-up time from baseline to CAD endpoint, death, or end of registry follow-up was 16.1 (0.1–20.1) years in EUFAM and 12.6 (0.02–19.0) years in FINRISK. More detailed information is given in the Supplementary material online.

### Lipidomics measurements

Lipidomic profiling and analysis of circulating lipid species was performed for 550 EUFAM family members with available plasma samples and for 897 individuals from the FINRISK 2012 cohort. Mass spectrometry-based lipid analysis was performed at Lipotype GmbH (Dresden, Germany) as described.^16^ Plasma and serum lipids were extracted with methyl tert-butyl ether/methanol (7:2, V:V) as in Matyash et al.^24^ Samples were analyzed by direct infusion in a QExactive mass spectrometer (Thermo Scientific) equipped with a TriVersa NanoMate ion source (Advion Biosciences). Samples were analyzed in both positive and negative ion modes in a single acquisition.

Data were analyzed with in-house developed lipid identification software based on LipidXplorer.^25, 26^ Reproducibility was assessed by the inclusion of reference plasma samples. Median coefficient of variation was <10% across all batches. A total of 151 species were detected in ≥ 80% of both EUFAM and FINRISK samples and were included in the subsequent analyses. Right-skewed lipidomics measures were natural logarithm transformed prior to normalization by standard deviation. More detailed information is given in the Supplementary material online.

### Statistical analyses

To assess the risk of incident coronary artery disease associated with the hyperlipidemias, we used Cox proportional hazards models using age as the time scale, stratified by sex, and clustered by family to estimate hazard ratios (HR) for incident CAD events, excluding individuals with prevalent CAD.

We used linear mixed models to estimate the association between lipidomic parameters and predictors of interest (hyperlipidemia status, continuous lipid measurement, or genotype) as implemented in MMM (version 1.01).^27^ Age, age^2^, and sex were used as additional fixed effect covariates. To account for relatedness among individuals, an empirical genetic relationship matrix was included as the covariance structure of a random effect. Statistical significance was evaluated using the Benjamini-Hochberg method at the 5% level to account for multiple comparisons. R (version 3.4.3) was used for data transformations and other analyses.^28^ Detailed information is given in the Supplementary material online.

## Methods

### Clinical characteristics and CAD risk of individuals with high levels of LDL-C or TGs

We first assessed the risk of developing CAD associated with high levels of LDL-C or TGs in individuals from the Finnish FINRISK population survey and in hyperlipidemic pedigrees ascertained as part of the European Study of Familial Hyperlipidemias (EUFAM) (Table 1; Supplemental Table I; Supplemental Figure I.A.). Individuals with high LDL-C had an increased risk of being diagnosed with CAD in the FINRISK population surveys (*n* = 19,644 individuals) compared to other individuals (HR 1.74; 95% CI 1.48–2.05). The members of hyperlipidemic families with high LDL-C had a similar though nonsignificant HR compared to their relatives without high LDL-C in 47 “high LDL-C” families (n = 633 individuals) (HR 1.71; 95% CI 0.94–3.10). The HRs did not differ between the cohorts (p=0.60). We also observed an increased CAD risk in individuals with high TGs in the population (HR 1.38; 95% CI 1.08–1.75), and a similar though nonsignificant HR in 35 “high TG” families (*n* = 375 individuals) (HR 1.35; 95% CI 0.52–3.51). The HRs did not differ between the cohorts (*p*=0.91). Meta-analyses of HRs closely approximated estimates derived in the population cohort.

**Table 1.** Coronary artery disease risk in familial and population-ascertained hyperlipidemias. *Clinical incident CAD event endpoints were defined as either myocardial infarction or coronary revascularization. Cox proportional hazards models, stratified by sex and clustered by family were used to estimate hazard ratios (HR) for incident CAD events using age as the time scale and excluding individuals with prevalent CAD*.

| Hyperlipidemia type | Parameter | Hyperlipidemic families | Population cohort | Meta-analysis |
| --- | --- | --- | --- | --- |
| High LDL-C | *n* (hyperlipidemic / non-hyperlipidemic) | 625 (136/489) | 19,644 (2,175/17,469) | 20,269 (2,311/17,958) |
|  | Number of incident CAD diagnoses (hyperlipidemic / non-hyperlipidemic) | 45 (16/29) | 904 (176/728) | 949 (192/757) |
|  | CAD HR (95% CI) | 1.71 (0.94-3.10) | 1.74 (1.48-2.05) | 1.74 (1.48-2.04) |
| High TGs | *n* (hyperlipidemic / non-hyperlipidemic) | 371 (72/299) | 19,644 (1,405/18,239) | 20,015 (1,477/18,538) |
|  | Number of incident CAD diagnoses (hyperlipidemic / non-hyperlipidemic) | 21 (3/18) | 904 (74/830) | 925 (77/848) |
|  | CAD HR (95% CI) | 1.35 (0.52-3.51) | 1.38 (1.08-1.75) | 1.38 (1.09-1.74) |

We then characterized the detailed lipidomic profiles of 550 individuals from 73 hyperlipidemic families and 897 individuals from the FINRISK population study (Methods; Supplemental Table II; Supplemental Figure I.B.). These included 105 individuals (23 %) out of 463 family members in 53 “high LDL-C” families who had LDL-C levels >90^th^ percentile (mean ± SE 5.2 ± 0.8 mmol/l), and 64 individuals (22 %) out of 287 family members in 39 “high TG” families who had TGs >90^th^ percentile (3.6 ± 2.0 mmol/l). Using similar cutoffs in the population, 56 individuals (6 %) and 65 individuals (7 %) out of 897 individuals were affected by high LDL-C levels (5.3 ± 1.1 mmol/l) and high TGs (3.5 ± 1.6 mmol/l) respectively. Simultaneously high LDL-C and TG levels were observed in 31 individuals in the family samples and 9 individuals in the population samples.

### High LDL-C and lipidomic profiles

To characterize the lipidomic profiles associated with elevated values of LDL-C, we compared individuals with high LDL-C levels vs. those without. In the hyperlipidemic families, individuals with high LDL-C had significantly elevated levels of 99 lipid species spread out across most of the studied lipid classes. Reduced levels were observed for three lysophospatidylcholine (LPC), two lysophosphatidylethanolamine (LPE) and one phosphatidylcholine-ether (PCO) species (Figure 1.A.; Supplemental Table III). Similar trends were seen in the population when comparing individuals with high LDL-C vs. those without. We observed significantly elevated levels of 51 lipid species (Figure 1.B.; Supplemental Table III). The effect estimates correlated strongly across all lipid species between the hyperlipidemic families and the population cohorts (Pearson’s r = 0.80; Figure 3). Further, we observed no statistically significant cohort-dyslipidemia interactions at the 5% false discovery rate.

**Figure 1.**
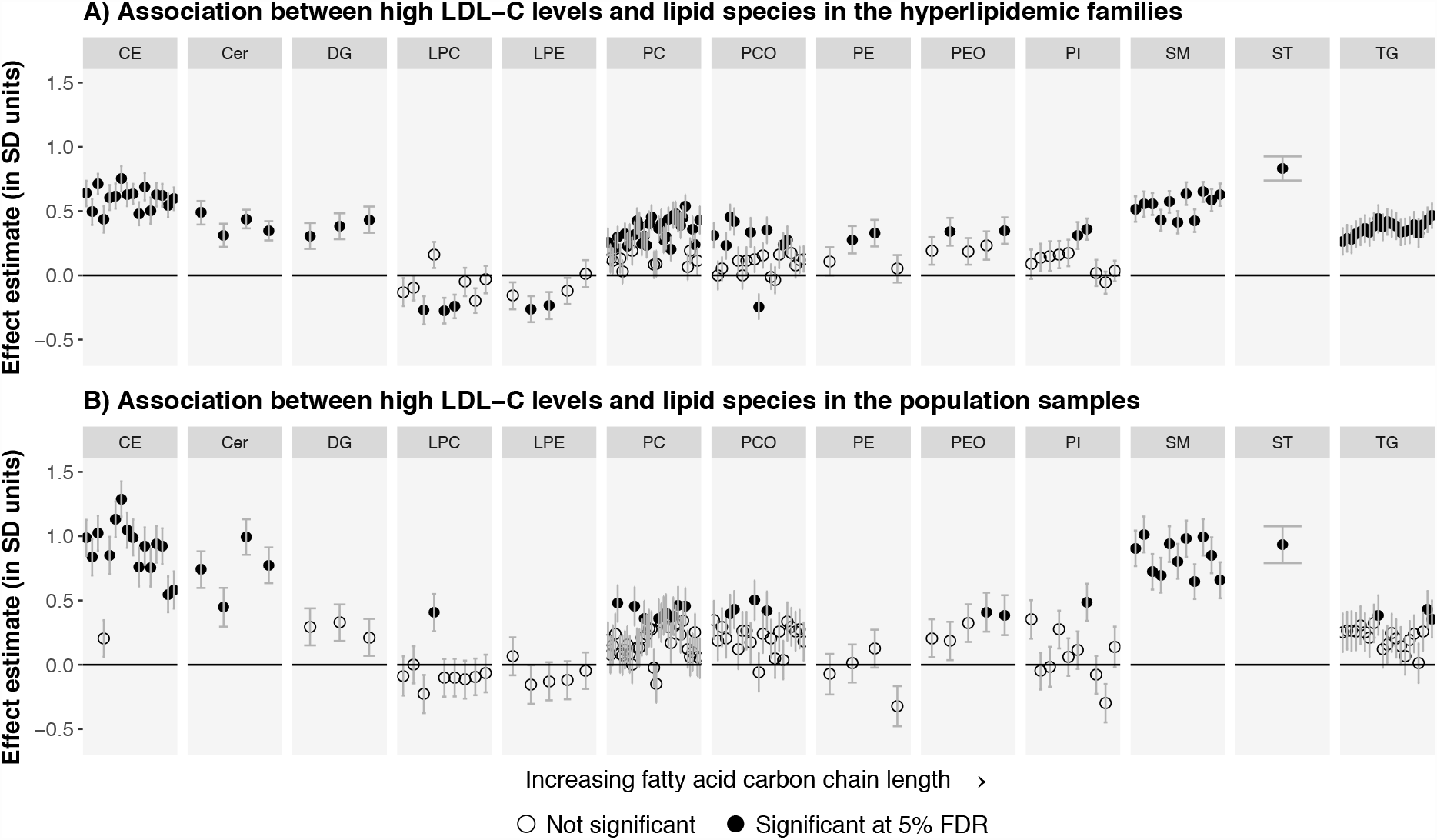
Associations between high LDL-C status and the levels of 151 lipid species. Panel A), individuals affected by high LDL-C levels (*n*=105) were compared with their unaffected relatives (*n*=358) in the 53 “high LDL-C” families. Panel B), individuals affected by high LDL-C (*n*=56) were compared with other individuals (*n*=841) in the FINRISK population cohort. The association of high LDL-C status with the lipid species was estimated using linear mixed models with age, age^2^, and sex as the other fixed effect covariates. Statistical significance was evaluated using the Benjamini Hochberg method at a 5% false discovery rate (FDR). The ordering of the lipid species within each class is the same as in Supplemental Table III. *Cer = ceramide, DG = diacylglyceride, LDL-C = low-density lipoprotein cholesterol, LPA = lysophosphatic acid, LPC = lysophosphatidylcholine, LPE = lysophosphatidylethanolamine, PC = phosphatidylcholine, PCO = phosphatidylcholine-ether, PE = phosphatidylethanolamine, PEO = phosphatidylethanolamine-ether, PI = phosphatidylinositol, CE = cholesteryl ester; SM = sphingomyelin, ST = sterol, TG = triacylglyceride*.

We also studied the association of high LDL-C levels with the degree of saturation of fatty acids in each lipid class. In the hyperlipidemic families, high LDL-C levels were associated with increased saturation of LPCs and ceramides, as well as reduced saturation of LPEs, PCs, phosphatidylcholine-ethers (PCOs), and phosphatidylinositols (PIs) (p-value range = 0.019–0.0014) (Supplemental Figure II). In the population samples, the trends were similar, although there was an association for increased LPC saturation only (*p*=7.2*10^−4^). There were no cohort-hyperlipidemia interactions in any of the lipid species at the 5% false discovery rate. Overall, the lipidomic profiles associated with high LDL-C levels appeared similar in the hyperlipidemic families and the general population.

### High TGs and lipidomic profiles

In the hyperlipidemic families, individuals with high TGs had elevated levels of 107 lipid species covering all studied lipid classes with the exception of LPEs. In addition, we observed reduced levels of seven PCO, two LPC, and one PI species (Figure 2.A.; Supplemental Table III). Similar profiles were seen in the population when comparing individuals with high TGs vs. those without. We observed elevated levels of 108 species and reduced levels of ten PCO and one LPC species (Figure 2.B.; Supplemental Table III). The effect estimates correlated very highly across all species between families and population samples (Pearson’s r = 0.96; Figure 3). Further, we observed no cohort-dyslipidemia interactions for any of the lipid species at the 5% false discovery rate.

**Figure 2.**
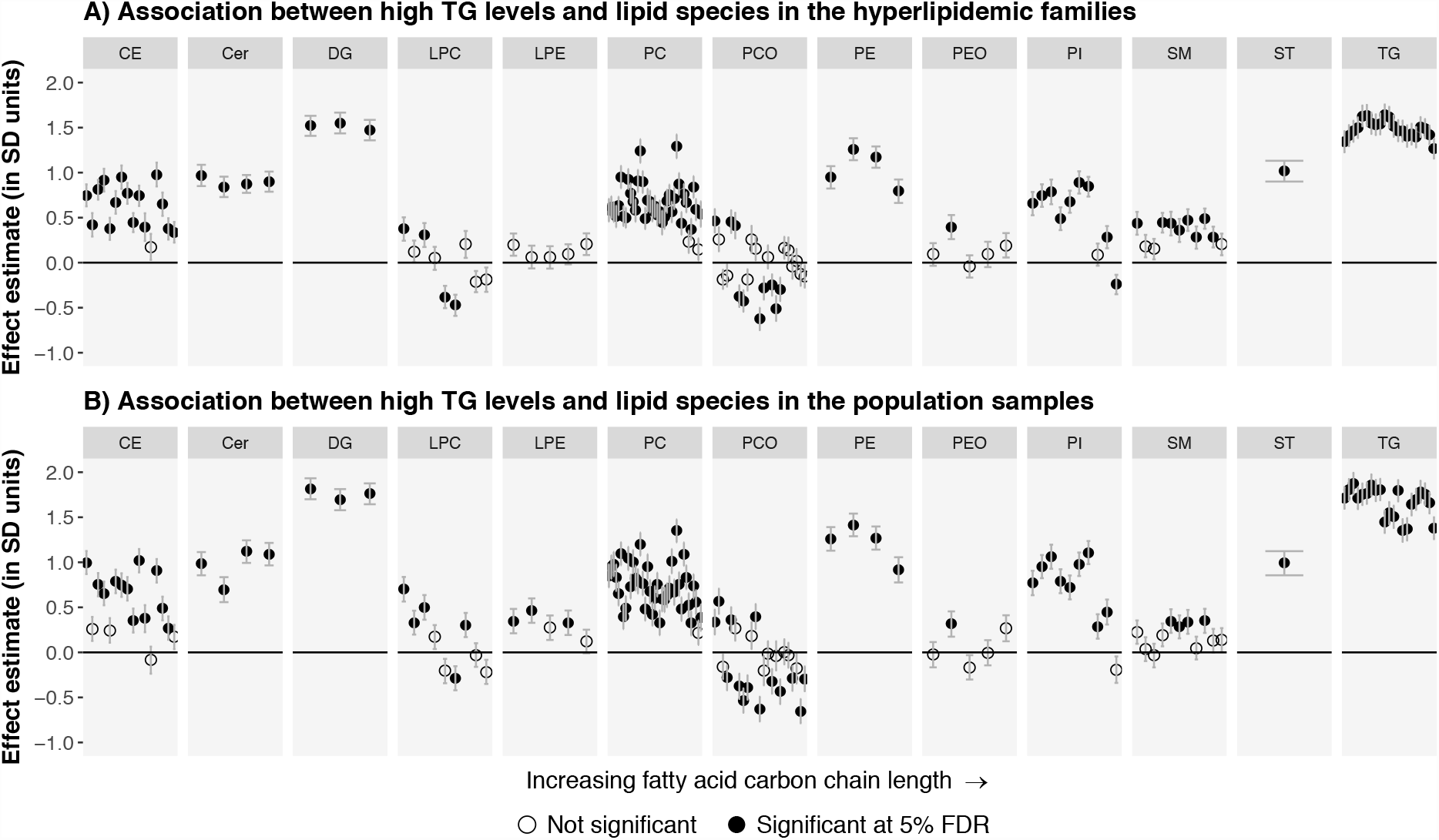
Associations between high TG status and the levels of 151 lipid species. Panel A), individuals affected by high TGs (*n*=64) were compared with their unaffected relatives (*n*=223) in 39 “high TG” families. Panel B), individuals affected by high TGs (*n*=65) were compared with other individuals (*n*=832) in the FINRISK population cohort. The association analyses were performed similarly toFigure 1. Abbreviations are displayed in Figure 1 legend.

**Figure 3.**
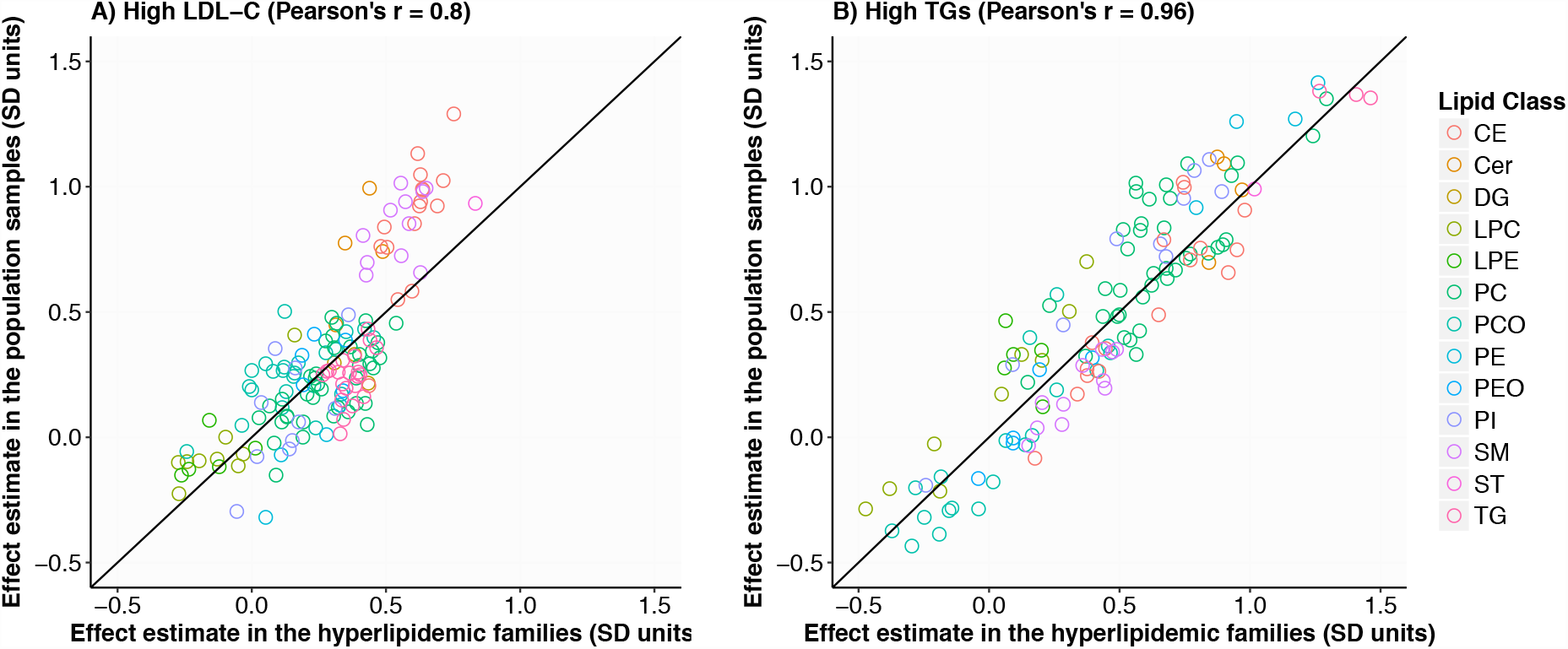
Correlation of effect estimates for hyperlipidemia status between the hyperlipidemic families and the population samples. The correlation between the effect estimates observed in the family and population cohorts is presented in A) for high LDL-C (effect estimates presented in Figure 1) and B) for high TGs (effect estimates presented in Figure 2). Abbreviations are displayed in Figure 1 legend.

Next, we studied the association of high TG levels with the degree of saturation of fatty acids in each lipid class. In both the hyperlipidemic families and the population, having high TGs was significantly associated with increased saturation of TGs, DGs, LPCs, and CEs (*p*-value range = 0.0012–5.9*10^−11^) (Supplemental Figure III). There were no statistically significant cohort-hyperlipidemia interactions at the 5% false discovery rate. Overall, we observed great similarity in the lipidomic profiles associated with high TGs in the hyperlipidemic families and in the general population.

### Independent associations of LDL-C and TG values with the lipid species

Since we observed modest combined hyperlipidemia in both cohorts, we estimated the independent associations of LDL-C and TGs with each lipid species in co-adjusted models (Figure 4; Supplemental Table III). In these analyses, many of the observed associations with LDL-C were greatly diluted in magnitude. LDL-C levels remained most strongly associated with CE, SM, ceramide, PC and PCO species in both cohorts. A total of 83 species in the hyperlipidemic families and 91 species in the population were independently associated with LDL-C at the 5% false discovery rate. In contrast, TGs remained strongly associated with a wide range of lipid species, including all individual TG species, DGs, PCs, PEs, PIs, ceramides and a subset of CEs in both cohorts. A total of 125 species in the hyperlipidemic families and 124 species in the population were independently associated with TGs at the 5% false discovery rate. Overall, only 13 species were uniquely associated with LDL-C in either cohort, whereas 42 species were uniquely associated with TGs (Supplemental Figure IV).

**Figure 4.**
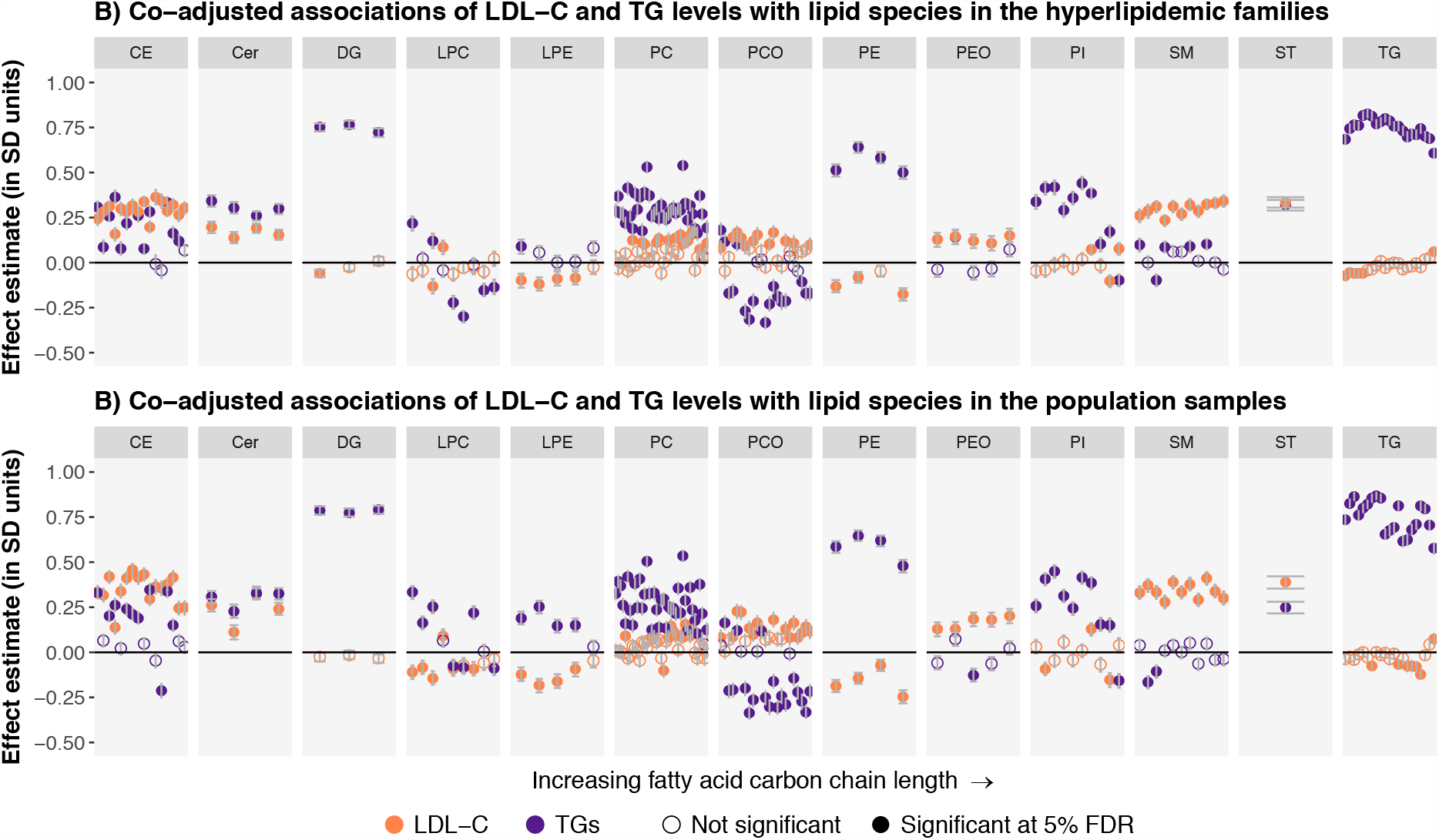
Independent (co-adjusted) associations of LDL-C and TGs with 151 lipid species. Effect estimates for LDL-C and TGs were derived from linear mixed models with the lipid species as outcomes, and LDL-C, log(TG^), age, age^2^, and sex as fixed effect covariates. The effect estimates were derived separately in the hyperlipidemic families (Panel A, *n* = 550 individuals) and the FINRISK population cohort (Panel B, *n* = 897 individuals). Effect estimates are presented for LDL-C in orange and TGs in purple. Statistical significance was evaluated using the Benjamini-Hochberg method at a 5% false discovery rate (FDR). The ordering of the lipid species within each class is the same as in Supplemental Table III. Abbreviations are displayed in Figure 1 legend.

## Discussion

Recent lipidomic approaches have identified several hundreds of different lipid species in the human circulation, some of which could be better prognostic biomarkers for atherosclerotic cardiovascular disease than the traditional clinical chemistry measurements. To the best of our knowledge, the present study is the most comprehensive lipidomic profiling of common hyperlipidemias to date. We used a mass spectrometric lipidomics platform to assess the lipidomic profiles in individuals with high LDL-C and/or TG levels. We found that individuals affected by high levels of LDL-C or TGs had CAD HRs between 1.34–1.74 in the family and population cohorts and exhibited distinct lipidomic profiles with clear variation between lipid classes. In total, out of 151 lipidomic species, 108 were significantly associated with high LDL-C and 131 with high TG levels in at least one cohort. Of these, 96 species were common with both high LDL-C and TGs. In addition, we observed highly similar lipidomic profiles between the familial and population-ascertained hyperlipidemias.

These findings allow us to draw several conclusions. First, the CAD risks are highly similar regardless of whether hyperlipidemic individuals were identified from families with a high prevalence of hyperlipidemia or from the general population. This result contrasts with earlier results reporting much higher CAD risks in relatives of familial combined hyperlipidemia probands compared to spouses.^29^ Our study, however, compares the estimates between family members to individuals with closely similar lipid levels from the population to isolate the effect of familiality. We also study the risk associated with elevated LDL-C and TGs separately. Our estimates are also lower than typically reported for monogenic FH, which may be associated with lifelong exposure to very high LDL-C levels and increased CAD risk compared with other individuals with similarly extreme hyperlipidemias.^8^ In the present study, we excluded probands with monogenic FH based on a functional LDL receptor test. Excepting monogenic FH, familial hyperlipidemias with high LDL-C and/or TG levels have been reported to be highly polygenic.^9, 10, 13, 30^ The pleiotropic effects of diverse genes, in contrast with the single affected pathway in monogenic FH, may partly explain why we did not observe increased CAD risk due to familiality in our study.

Second, to more deeply characterize potential differences between familial and population ascertained hyperlipidemias, we performed precise phenotyping of circulating lipid species known to be associated with ASCVD risk.^20–22^ We first characterized the lipid profiles associated with high LDL-C and TG levels. Many of the associations were not specific to LDL-C but rather due to combined dyslipidemia. LDL particles are generated in circulation as downstream metabolic products from the TG-enriched lipoproteins (liver-derived VLDL particles and their post-lipolytic remnants) by the action of two lipases, lipoprotein lipase and hepatic lipase.^31,32^ Thus, a proportion of the core lipids—especially cholesterol esters—and the particle surface phospholipids are retained within the generated LDL particles. The actions of the cholesteryl ester transfer protein and phospholipid transfer protein further modulate the constituents of TG-rich and LDL particles.^33^ Percentual lipid compositions have been reported for different lipoprotein classes, but they do not directly reflect variation in plasma LDL-C or TG concentrations. For example, PCs have been estimated to constitute 12 % of all lipids in LDL particles vs. 3–9% in TG-rich lipoproteins.^34^ However, in our study, PCs were overall more strongly associated with TG levels than with LDL-C levels. Nevertheless, LDL-C remained positively associated with a range of species including CEs, ceramides, SMs, PCs and PCOs. Among the strongly increased species, CE(14:0), CE(16:0), CE(16:1), CE(18:0), SM(34:1;2), SM(34:2;2), SM(42:2;2), Cer(42:1;2), and Cer(42:2;2) have previously been associated with the risk of future ASCVD incidence or events.^20^,

Elevated TG levels were associated with either increases or decreases in the levels of lipid species across most classes. Importantly, most of these associations appeared to be independent of LDL-C levels. Among the lipid species that were strongly correlated with high TGs even after correction for LDL-C levels were several species which have previously been associated with risk of ASCVD incidence or events.^20–22^ These include the species CE(14:0), CE(16:0), CE(16:1), CE(18:0), TG(50:1), TG(50:2), TG(50:3), TG(52:2), TG(52:3), TG(52:5), TG(56:5), TG(56:6), Cer(42:1;2) and Cer(42:2;2). Further, high TGs were associated with increased saturation of fatty acids in the TG, DG, CE and LPC classes. Such differences in the relative concentrations of circulating fatty acids can be partly related to dietary intake, but are also influenced by endogenous metabolism.^35^

Overall, a larger proportion of the lipid species previously linked with increased ASCVD risk were strongly associated with elevated TGs rather than with elevated LDL-C. This interesting observation suggests that the levels of these lipid biomarkers are more closely linked with circulating TG-rich lipoprotein metabolism than with low-density lipoproteins.

Third, several lipid species, such as specific cholesteryl esters, ceramides, and phosphatidylcholine-ethers, remained independently associated with both elevated LDL-C and TGs. Among these species, the ceramides Cer(42:1;2) (presumably Cer(d18:1/24:0)) and Cer(42:2;2) (presumably Cer(d18:1/24:1)), the sterol esters CE(16:1) and CE(18:0), and the sphingomyelin SM(34:1;2) may have added value in the prediction of ASCVD incidence or event risk over traditional lipid measurements.^20–22^ Plasma ceramides have been reported to be independent predictors of cardiovascular events in addition to LDL-C in the population and in CAD patients.^22, 36, 37^ Both LDL-C and TGs remained independently associated with all four ceramides quantified in our study, and LDL-C was additionally associated with increased saturation degree of ceramides. Unlike most CE species, CE(16:1) was more strongly associated with the concentration of TGs than with LDL-C concentration in our study. SM(34:1;2) was the only sphingomyelin species which was negatively associated with TGs – and this association became evident only after adjusting for LDL-C levels. Additionally, some species, such as Cer(42:1;2) and TG(56:6), which were positively associated with hyperlipidemias in our sample, have previously been reported to be associated with decreased risk of ASCVD events.^20, 22^ These co-associations and discordances between reported associations might explain why some lipid species can improve risk prediction. Consequently, there is an urgent need for a better understanding of the potential underlying signaling and metabolic pathways.

Finally, the lipidomic profiles associated with high LDL-C or TG levels were comparable between familial and population-ascertained hyperlipidemias. We observed no differences in either the levels of individual lipid species or the saturation of fatty acids within lipid classes. Our results support the hypothesis that familial and population-ascertained hyperlipidemias have similar, overlapping, and heterogeneous pathophysiology. Our results are also reassuring for the quality of lipidomics studies which combine familial and population-based hyperlipidemic samples to increase statistical power.

Although we present the most comprehensive characterization of coronary artery disease risk and circulating lipid species in common familial hyperlipidemias to date, our study has some limitations.

The confidence intervals for CAD risk were relatively large in the family samples. The field of lipidomics is still relatively young, and concerns have been raised regarding the replicability of individual lipidomics platforms. The platform used here overcomes these problems by profiting from the advantages of direct infusion mass spectrometry for high throughput screening studies. We also observed similar lipidomic profiles associated with hyperlipidemia in two independent cohorts, supporting the replicability of the platform. Further, the lipid species included in our analyses are strongly associated with known genetic lipid loci (Supplemental Figures V-VIII). We profiled the whole circulating lipidome at once to capture variation across various lipoprotein classes and size distributions.

However, this limits our ability to follow up on observed differences at the level of the individual lipoproteins. We excluded poorly captured lipid species from the analyses; future advances in lipidomics technology might enable their detection. It is unclear how well our results can be generalized to other populations than Finns. Some of the individuals surveyed in population cohorts might in fact have familial hyperlipidemia; we could not fully rule out such cases. As this was a cross-sectional study, single LDL-C and TG measurements were used to establish hypercholesterolemia status. However, especially TGs are known to exhibit diurnal variability—especially in relation to dietary episodes with chylomicron entry—as well as variation over time periods of months to years.^38–40^ In this study, the samples from the hyperlipidemic families were fasting and the samples from the population cohort were semi-fasting. In this light, the similarity of lipidomic profiles between familial and population ascertained hypertriglyceridemias becomes even more striking. Moreover, the most recent consensus statement from the European Atherosclerosis Society and European Federation of Clinical Chemistry and Laboratory Medicine recommends routine use of non-fasting blood samples for assessment of plasma lipid profiles.^41^

In conclusion, we observed distinct lipidomic profiles associated with high levels of LDL-C and TGs, but similar CAD risk and lipidomic profiles between common familial and population-ascertained hyperlipidemias, providing further evidence of similar and overlapping underlying mechanisms. The high degree of similarity between these familial dyslipidemic individuals and random population samples in their genomic and lipidomic profiles, and in elevated CAD risk, does not support different screening and treatment for familial and sporadic cases. Additional work is needed to confirm the validity of this hypothesis in clinical settings.

## Acknowledgements

We would like to thank Sari Kivikko, Huei-Yi Shen, and Ulla Tuomainen for management assistance. The FINRISK data used for the research were obtained from THL Biobank. We thank all study participants for their generous participation in the FINRISK and EUFAM studies.

